# The Batten disease protein CLN3 is important for stress granules dynamics and translational activity

**DOI:** 10.1101/2022.05.20.492784

**Authors:** Emily L. Relton, Nicolas J. Roth, Seda Yasa, Abuzar Kaleem, Guido Hermey, Stephane Lefrancois, Peter J. McCormick, Nicolas Locker

## Abstract

The assembly of membrane-less organelles such as stress granules (SGs) is emerging as central in helping cells rapidly respond and adapt to stress. Following stress sensing, the resulting global translational shutoff leads to the condensation of stalled mRNAs and proteins into SGs. By reorganising cytoplasmic contents, SGs can modulate RNA translation, biochemical reactions and signalling cascades to promote survival until the stress is resolved. While mechanisms for SG disassembly are not widely understood, the resolution of SGs is important for maintaining cell viability and protein homeostasis. Mutations that lead to persistent of aberrant SGs are increasingly associated with neuropathology and a hallmark of several neurodegenerative diseases. Mutations in *CLN3* are causative of juvenile neuronal ceroid lipofuscinosis (JNCL), a rare neurodegenerative disease affecting children. *CLN3* encodes a transmembrane lysosomal protein implicated in autophagy, endosomal trafficking, metabolism, and response to oxidative stress. Using a HeLa KO model, we now show that CLN*3*^KO^ is associated with an altered metabolic profile, reduced global translation, and altered stress signalling. We further demonstrate that loss of CLN3 results in perturbations in SG dynamics, resulting in assembly and disassembly defects, and altered expression of the key SG nucleating factor G3BP1. With a growing interest in SG-modulating drugs for the treatment of neurodegenerative diseases, novel insights into the molecular basis of CLN3 Batten disease may reveal avenues for disease-modifying treatments for this debilitating childhood disease.

## Introduction

Neuronal ceroid lipofuscinoses (NCL) are a group of fatal inherited neurodegenerative diseases that affects all age groups, but is predominantly diagnosed in children. Mutations in one of 13 distinct genes (*CLN1 - CLN8, CLN10 - CLN14*) are the cause of NCL (1). The most common form of NCL is CLN3 disease, also known as Batten disease. Symptoms include visual loss, cognitive decline, ataxia and seizures. Symptoms most often appear between the ages of 5 and 8 years, with death occurring in the 3rd decade of life (2). The most common mutation found in patients is the deletion of exon 7 and 8, resulting is a significantly truncated protein, but other less frequent mutations have also been identified (1,3).

CLN3 is a highly glycosylated integral membrane protein with its N- and C-terminal ends located in the cytosol. The protein is localized to various intracellular compartments including endolysosomes (4), and has proposed roles in intracellular trafficking and autophagy (5-7). CLN3 can interact with Rab7A (8), a small GTPase that regulates the spatiotemporal recruitment of retromer (9,10). Retromer is a protein complex that is required for the endosome-to-TGN retrieval of the lysosomal sorting receptors, mannose 6-phosphate receptor and sortilin (11,12). CLN3 can interact with retromer and the sorting receptors, and functions as a scaffold protein to ensure efficient interactions (13). In cells lacking CLN3, retromer does not interact with the sorting receptor, which results in their lysosomal degradation (15). This results in inefficient sorting of soluble lysosomal proteins, leading to defective lysosomes. Autophagy defects found in CLN3-deficient cells could be due to the lack of efficient lysosomal protein sorting, or could be due to the lack of fusion of autophagosome with lysosomes (14), or a combination of mechanisms.

Due to their highly specialised nature, neurons have an important energy demand, which is met by mitochondrial production of large quantities of ATP by oxidative phosphorylation. Therefore, neurons are especially vulnerable to defects in homeostatic equilibrium and metabolic stresses. Previous studies have suggested that CLN3 is important for maintaining metabolic homeostasis. The loss of CLN3 is associated with the downregulation of oxidative phosphorylation and glycolytic enzymes, and reduced glycolytic metabolites, suggesting impaired glycolysis (15,16). Moreover, mammalian models of CLN3 disease display altered expression of mitochondrial enzymes involved in cellular respiration, increased accumulation of reactive oxygen species and membrane depolarisation linked to higher cell death levels (17). Studies in drosophila have also revealed a hyper-sensitivity to oxidative insult for CLN3 mutants (18). Finally, observations in patients’ fibroblasts have led to the proposal that ROS accumulation is a common hallmark of NCLs and may be correlated with disease severity (19). In addition to these metabolic defects, CLN3 has also been linked to the resolution of ER stress and its resolution via the unfolded protein response (UPR). The overexpression of wild-type CLN3 appears to be protective against tunicamycin induced ER stress, increasing expression of ER chaperone glucose-related protein 78 (GRP78), and reduced expression of apoptotic markers such as C/EBP homologous protein (CHOP). In contrast, depletion of CLN3 produced the opposite effect, suggesting its involvement in resolving ER stress (20).

The reorganisation of cellular content into stress granules (SGs) is a prime feature of the cellular response to many stresses including viral infection, protein misfolding, nutrient depletion, UV, heat and oxidative stress. In response to these stresses, the general inhibition of protein synthesis results in the dissociation of mRNAs from polysomes and their accumulation in RNP complexes (21). This increased concentration of cytoplasmic RNPs and their binding by aggregation prone RNA-binding proteins (RBPs), such as Ras-GTPase activating SH3 domain binding protein 1 (G3BP1), results in the recruitment of multiple resident proteins characterised by the presence of low sequence complexity, and intrinsically disordered regions in their structure. These drive clustering/fusion events supported by multivalent interactions between their protein and RNA components, with G3BP1 acting as a key node for promoting RNA-protein, protein-protein and RNA-RNA interactions, ultimately resulting in liquid-liquid phase separation (LLPS) and SG formation (22,23). SGs are highly dynamic, rapidly assembling to sequester the bulk content of cytoplasmic mRNAs, and dissolving upon stress resolution to release stored mRNAs for future translation (24,25).

By sequestering specific proteins following stress, SGs alter the composition and concentration of cytoplasmic proteins, and therefore have been proposed to modulate biochemical reactions and signalling cascades in the cytosol, impacting on signalling and metabolism (26-28). Importantly, many signalling molecules associated with diseases can concentrate in SGs, and several are regulated by SGs, supporting a role for SGs as signalling hubs (29). However, because of their proposed stress-specific nature and composition, the exact role for SGs in different stresses remains poorly understood and while protective functions have been ascribed to SGs, they are also suggested to support pro-death functions (30). SGs are dynamic and reversible, with their disassembly important to restore translation and maintaining protein homeostasis after stress. Mutations impacting protein turnover, or in SG resident RNA-binding proteins (RBPs) impeding SG clearance or dysregulating LLPS, lead to persistent or aberrant SGs, are emerging as a determinant of neuropathology, in particular in amyotrophic lateral sclerosis (ALS) and related diseases (31). For example, ALS causing mutations cluster in low-complexity RG-rich regions of the RBP TDP43 or FUS, and alter phase transition properties and dynamics of SGs (31,32). In addition, non-favourable conditions within the disease micro-environment can impair the cells ability to respond to additional stressors resulting in the promotion of apoptosis (32). Finally oxidative cell environments, caused by the accumulation of ROS, inhibit SGs via the oxidation of core SG proteins involved in their nucleation (33). Collectively these data provide clear evidence linking the pathophysiology of neurodegenerative diseases with SG biology.

Because of the previous studies linking CLN3 with metabolic stress discussed above, and with further studies linking CLN3 and the resolution of endoplasmic reticulum (ER) stress, we set out to investigate the unexplored interplay between CLN3 Batten disease and the SG pathway. Using CLN3 knock out (CLN3^KO^) models, we demonstrate that the loss of CLN3 is associated with a defect in mitochondrial activity in HeLa cells, the rewiring of signalling pathways controlling translation and a reduction of protein synthesis. In turn, CLN3 is important for the assembly of SGs and their resolution upon stress removal, with the accumulation of persistent SGs occurring in CLN3^KO^ cells. This defect in SG assembly/disassembly dynamics are not associated with impaired stress signalling via the integrated stress response or a reduced ability of SG resident proteins to phase separate. Rather, we propose that reduced levels of G3BP1 in CLN3^KO^ cells are responsible for these defects. These results shed light for the first time on the importance of CLN3 in SG biogenesis and given the recent interest in SGs as druggable targets may provide novel therapeutic opportunities for a rare juvenile disease with currently no efficient treatments.

## Results

### CLN3^KO^ cells display impaired basal metabolic function and are hyper-sensitive to additional stress

Previous studies have shown that the loss of CLN3 is associated with perturbations in metabolic homeostasis (15,16). We first tested if this could be observed in our cell model by comparing the metabolic phenotypes of wild-type (WT) and CLN3^KO^ HeLa cells. The rates of oxidative phosphorylation and glycolysis was determined by measuring the oxygen consumption rate (OCR) and extracellular acidification rate (ECAR) shown in fig1A and fig1B, respectively. CLN3^KO^ cells displayed significantly lower OCR and ECAR, suggesting that CLN3^KO^ lowers the rates of both glycolysis and mitochondrial respiration by oxidative phosphorylation. Moreover, after the addition of the metabolic stressors oligomycin and FCCP, the difference in the rates of metabolism in the CLN3^KO^ compared to WT HeLa cells was more profound. These data suggest that CLN3^KO^ cells have an impaired ability to handle additional stressors and start with a lower metabolic turnover in the absence of stressors, which is indicative of impaired mitochondrial function.

**Figure 1:**
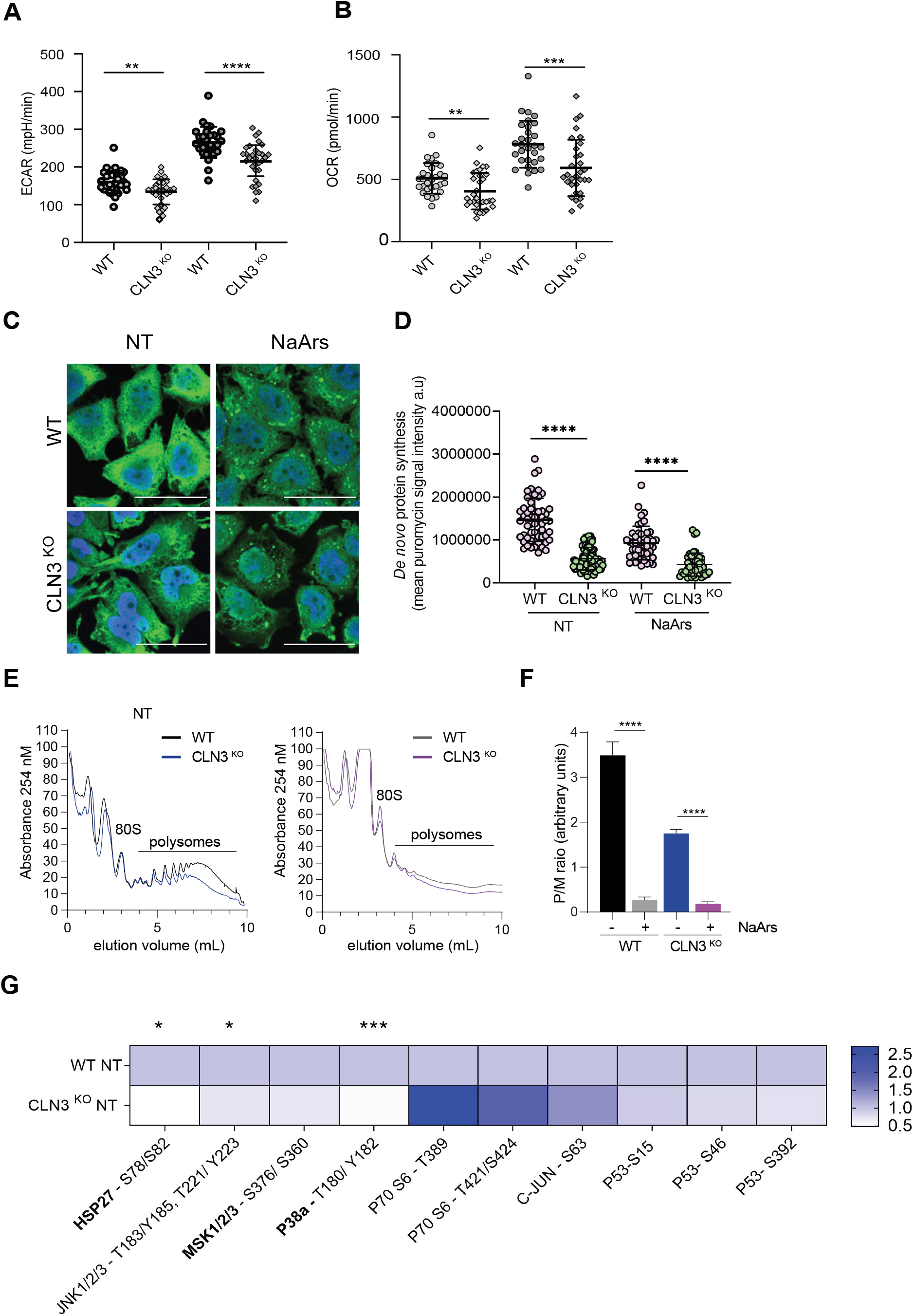
Metabolic, proteostatic and signalling defects accompany CLN3^KO^ HeLa cells. (A) Efficacy of glycolysis was measured by extracellular acidification rate (ECAR). CLN3^KO^ HeLa cells display reduced glycolytic flux as compared to WT at baseline conditions. The same trend is observed after challenge with NaArs which increases glycolytic flux irrespective of genotype. (B) Mitochondrial respiration was assessed using oxygen consumption rate (OCR). CLN3^KO^ HeLa cells display reduced OCR under NT (baseline) conditions. Challenge with NaArs increases OCR in both genotypes but is significantly lower in CLN3^KO^ HeLa cells (C) Immunofluorescent images of puromycin incorporation (green) with nuclei stained with dapi (blue). Puromycin signal intensity is correlated with translation efficacy in non-treated (NT) and NaArs treated cells. Scale bar = 40μm^2^. (D) Efficacy of *de novo* protein synthesis was assessed by measuring integrated density values of single cells. CLN3 KO displays significantly reduced translation in NT conditions compared to WT. One representative replicate is shown. (Mann-Whitney test p<0.0001). NaArs treatment exacerbates translational shut off in CLN3 KO compared to WT (Mann-Whitney test p<0.0001). (E) Polysome fractions in WT and CLN3^KO^ HeLa cells in NT condition. CLN3^KO^ HeLa cells exhibits fewer polysomes compared to WT in NT conditions characteristic of reduced rate of translation while NaArs treatment depletes polysome fraction in WT and CLN3^KO^ HeLa cells. HeLa cells were prepared as detailed under “Experimental Procedures”. A normalized (by A254) amount of lysate was separated on a 10–50% sucrose gradient and fractionated into 1-ml fractions using a FoxyR1 collection system (Teledyne). The displayed trace represents absorbance at 254nm (vertical axis) throughout the gradient from top (left) to bottom (right). 80 S (monosome), and polysome peaks are labeled. (F) The areas below the monosome and polysome peaks were determined for several biological replicates (n=3), and the mean polysome/monosome (P/M) ratio was calculated by measuring the area under the polysomal (P) to monosomal (M) peaks using standard area under the curve methods. ****, p<0.0001. (G) Heatmap shows differences in phosphorylation status of a panel of target residues spanning various stress-activated signalling pathways. Decreased phosphorylation of HSP27 (S78/S82), JNK1/2/3 (T183/Y185, T221/Y223) and p38α (T180/Y182). Results analysed by multiple t-tests (* p<0.02, *** p<0.0002).

### CLN3^KO^ display hallmarks of impaired protein synthesis

An increasing number of studies support a link between mitochondrial function and proteostasis failure, both defining characteristics of neurodegeneration (34). Given the observed differences in mitochondrial function, we hypothesised that potential alterations in bioenergetics would be deleterious to bulk protein synthesis, a vastly energy expensive process. To probe the impact of CLN3^KO^ on translational control, we assessed global translational efficiency using single cell analysis by measuring the incorporation of puromycin, a tRNA structural homolog which specifically labels actively translating nascent polypeptides and causes their release from ribosomes (35). Anti-puromycin antibodies are then used to detect puromycylated native peptide chains by confocal microscopy (Fig1C). Quantification of puromycin signal intensity revealed a significant reduction in *de novo* protein synthesis in CLN3^KO^ compared to WT cells under baseline conditions (Fig1D). To confirm these results and obtain a deeper understanding of the translational control associated with CLN3^KO^, we used polysome profiling to separate actively translating ribosomes in cytoplasmic extracts. Cell lysate from WT or CLN3^KO^ HeLa cells were prepared and subjected to 15-50% sucrose density centrifugation. A typical polysome profile pattern was obtained as shown on Fig1E. As expected, stimulation with Sodium Arsenite (NaArs) resulted in a total shut-down of translation and disappearance of polysomes (Fig1E). Furthermore, in contrast to WT cells, CLN3^KO^ cells exhibited a translational defect as shown by a decrease in the amount of polysomes in basal conditions. This was further quantified by monitoring the ratio of polysomes to monosomes. Quantification of the area under the polysome and monosome peak showed that the Polysome/Monosome (P/M) ratio, reflecting the amount of ongoing translation and mRNAs associated with translating ribosomes, was significantly reduced in CLN3^KO^ when compared to WT cells (Fig1F).

### CLN3^KO^ is associated with impaired stress-response signalling

Stress-activated signalling pathways sense perturbations in cellular stoichiometry and drive adaptation to maintain homeostasis. Breakdown in this complex signalling network between organelles is thought to contribute towards the common multifactorial pathologies associated with different neurodegenerative disease (36). We probed the phosphorylation status of effector proteins across several major stress-activated signalling pathways in response to the loss of CLN3 under basal conditions. Our results show differences in the phosphorylation of several stress activated kinases (Fig1G). These include significant decreases in phosphorylation at residues on heat shock protein 27 (S78/S82), JNK 1/2/3 (T183/Y185/T221/Y223) and P38α (T180/Y182), as well as non-significant yet trending increased phosphorylation of P70 S6 (T389/T421/S424). Together this data suggests that loss of CLN3 may hamper the stress signalling networks involved in sensing and responding to external and internal stressors.

### Stress granule assembly is perturbed by loss of CLN3

The link between SGs and the pathogenesis of neurodegeneration is well established (31), however the involvement of SGs in CLN3 disease has not been explored. SGs are important in determining cell fate in response to stressful stimuli, preventing apoptosis by rewiring the stress-activated signalling network (29,33). Because we observed differences in stress signalling following loss of CLN3, we investigated the ability of CLN3^KO^ HeLa cells to mount a SG response following acute oxidative insult. NaArs was used to induce SGs over a time course ranging from 5 to 45 mins. Cells were then fixed and labelled with anti-G3BP1 to detect SG assembly (Fig2A). First, we confirmed that G3BP1 colocalised with an additional SG marker, eIF3B, suggesting the proper assembly of NaArs-induced canonical SGs in our model cells (Sup Fig1). The percentage of cells displaying SGs was quantified using G3BP1 foci as an SG marker and SGs were categorised as small (0.1-0.75μm2), medium (0.75-6μm2) or large (>6μm2) (Fig2B). Small and medium SGs assembled from 5mins NaArs treatment and increased with length of NaArs exposure before reaching a peak at 20 mins of NaArs treatment. Large granules were observed from 20 mins, likely due to fusion of smaller granules. We observed 50% fewer large SGs at 20 min and 45 mins in CLN3^KO^ when compared to WT HeLa cells, suggesting an impairment in the maturation of SGs in the later stages of SG induction.

**Figure 2:**
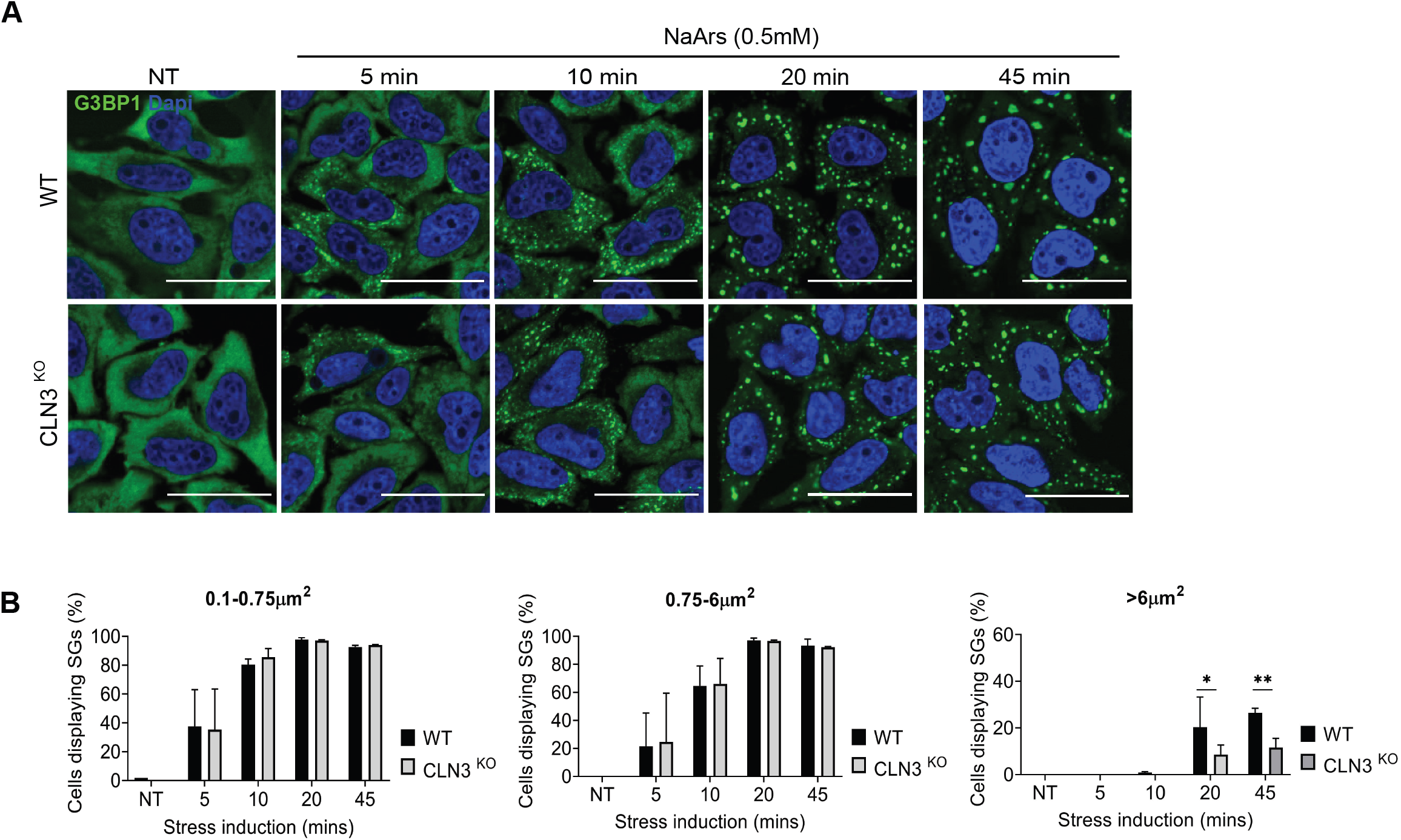
CLN3 KO displays defect in SG induction pathway. (A) Representative immunofluorescent images show time course of SG induction after NaArs treatment. Cells stained with G3BP1 green) and nuclei dapi stained (blue). Scale bar = 40μm^2^. (B) Quantification of small (0.1-0.75 μm^2^), medium (0.75-6 μm^2^) and large (>6 μm^2^) SGs shown as %cells displaying SGs. Results shown mean ± SD, n=3. CLN3^KO^ HeLa cells display significantly fewer large SGs after 20 (P=0.01) and 45 minutes (p=0.002) of NaArs induction. Quantification performed using Image J plugin Aggrecount. Statistical analysis was carried out using multiple t-tests.

### Aberrant stress granules disassembly dynamics is associated with loss of CLN3, but not CLN5

Abnormal turnover of SGs, in particular the persistence of SGs after stress is resolved, is now a well-established hallmark of neurodegenerative disease (31). To assess the impact of CLN3^KO^ on SG clearance, cells were stressed with 0.5mM NaArs for 45 mins to induce SGs. The stressor was subsequently washed out and cells were incubated with fresh media for a maximum of 3 hours. SGs were detected by immunofluorescence using G3BP1 and eIF3b as markers, as shown in Fig3A, and categorised as small, medium or large as previously described. The frequency of cells containing SGs was than quantified in an automated and non-biased manner using Aggrecount (37). As expected, NaArs treatment robustly induced the assembly of small and medium SGs which decreased in frequency in a temporally dependent manner after the removal of the stressor (Fig3B). At 3 hours post stress, the frequency of small and medium SGs had returned to non-treated (NT) levels in WT cells, however CLN3^KO^ cells displayed altered recovery dynamics. We observed that 35% of cells had persistent small SGs and 24% of cells contained medium sized SGs after 3 hours of recovery. This data suggests the clearance of NaArs induced SGs is less efficient in CLN3^KO^ and that CLN3 may be important for recovery and adaptation to oxidative stress.

**Figure 3:**
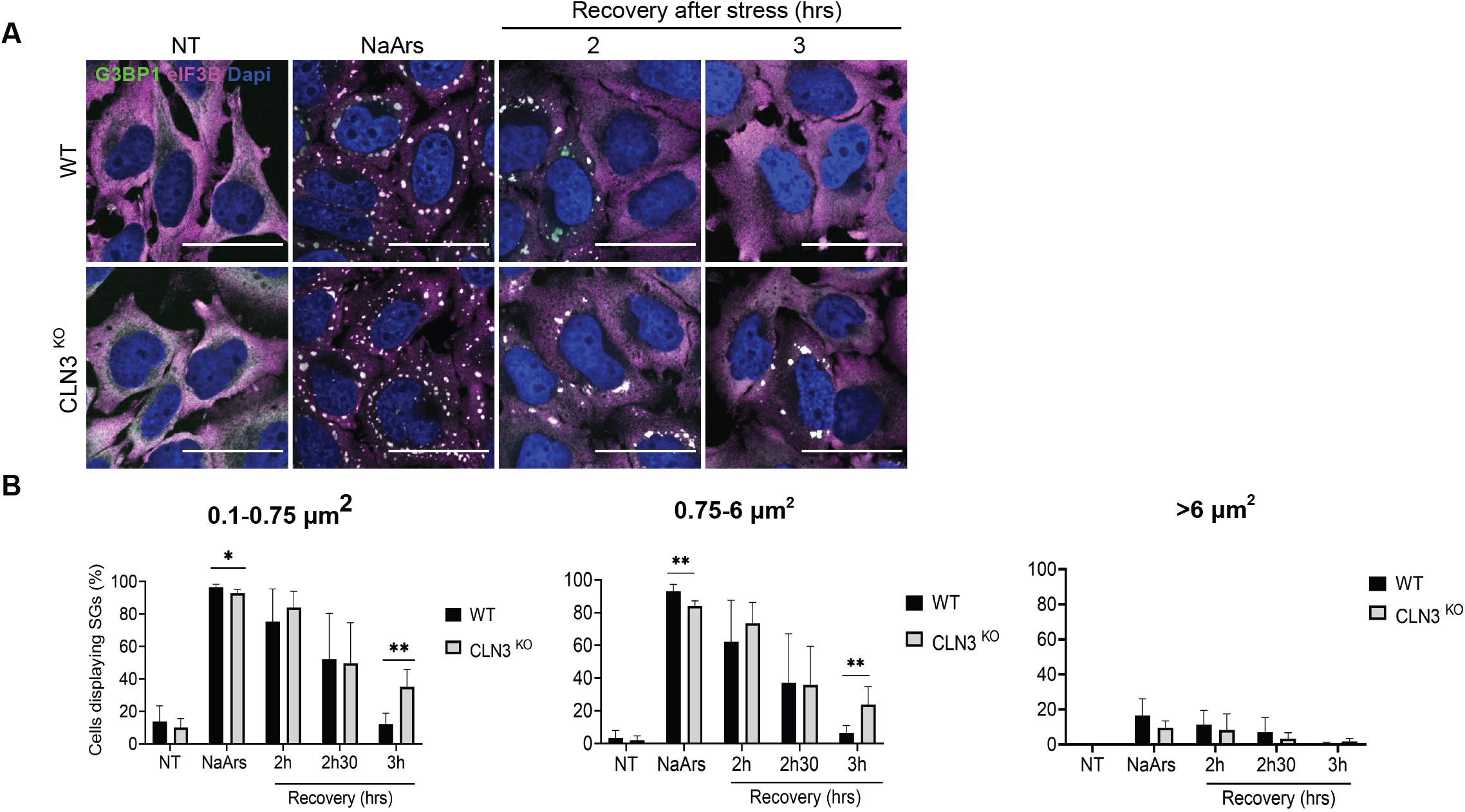
CLN3^KO^ HeLa cells display defect in clearance of NaArs-induced SGs. (A) Representative immunofluorescent images show time course of SG clearance after NaArs challenge. Scale bar = 40μm^2^. Cells stained for SG markers G3BP1 (Green) and Eif3b (Magenta). Nuclei stained with dapi (blue) (B) Quantification of small (0.1-0.75 μm^2^), medium (0.75-6 μm^2^) and large (>6 μm^2^) shown as %Cells displaying SGs. Results show mean ± SD, n=6. CLN3^KO^ HeLa cells display a significant number of persistent small (P=0.001) and medium SGs (P=0.005) at 3h recovery post stress compared to WT. Quantification was performed using Image J plugin Aggrecount. Statistical analysis was carried out using multiple t-tests.

Autophagy pathways have previously been linked to SG clearance (38). Furthermore, it has been previously shown that CLN3 is required for the endosome-associated small GTPAse RAB7A functions, including its interaction with PLEKHM1, an interaction required for endosome-lysosome and autophagosome-lysosome fusion events, as well as the efficient recycling of lysosomal receptors sortilin and CI-MPR (13) Therefore, we asked if impaired SG clearance was due to the loss of this CLN3-Rab7A interaction, however we found SG clearance was unaffected by Rab7A^KO^ HeLa cells (Sup Fig2). This suggests that the increased persistence of SGs in CLN3^KO^ is not due to CLN3-dependent Rab7A functions associated with lysosomal function.

NCL pathologies are linked to the dysfunction of several CLN proteins involved in intracellular trafficking (13,39). To assess if the delay in SG dissolution is CLN3 specific, we investigated SG dynamics in a HeLa KO model of CLN5 disease. NaArs induced small and medium SGs in 97% and 92% of CLN5^KO^ HeLa cells respectively. After wash out of the stressor and a 3hr recovery period we observed that the frequency of all sizes of SGs was comparable between WT and CLN5^KO^ HeLa cells, suggesting the loss of CLN5 does not impact the clearance of SGs and impaired recovery (Fig 4B).

**Figure 4:**
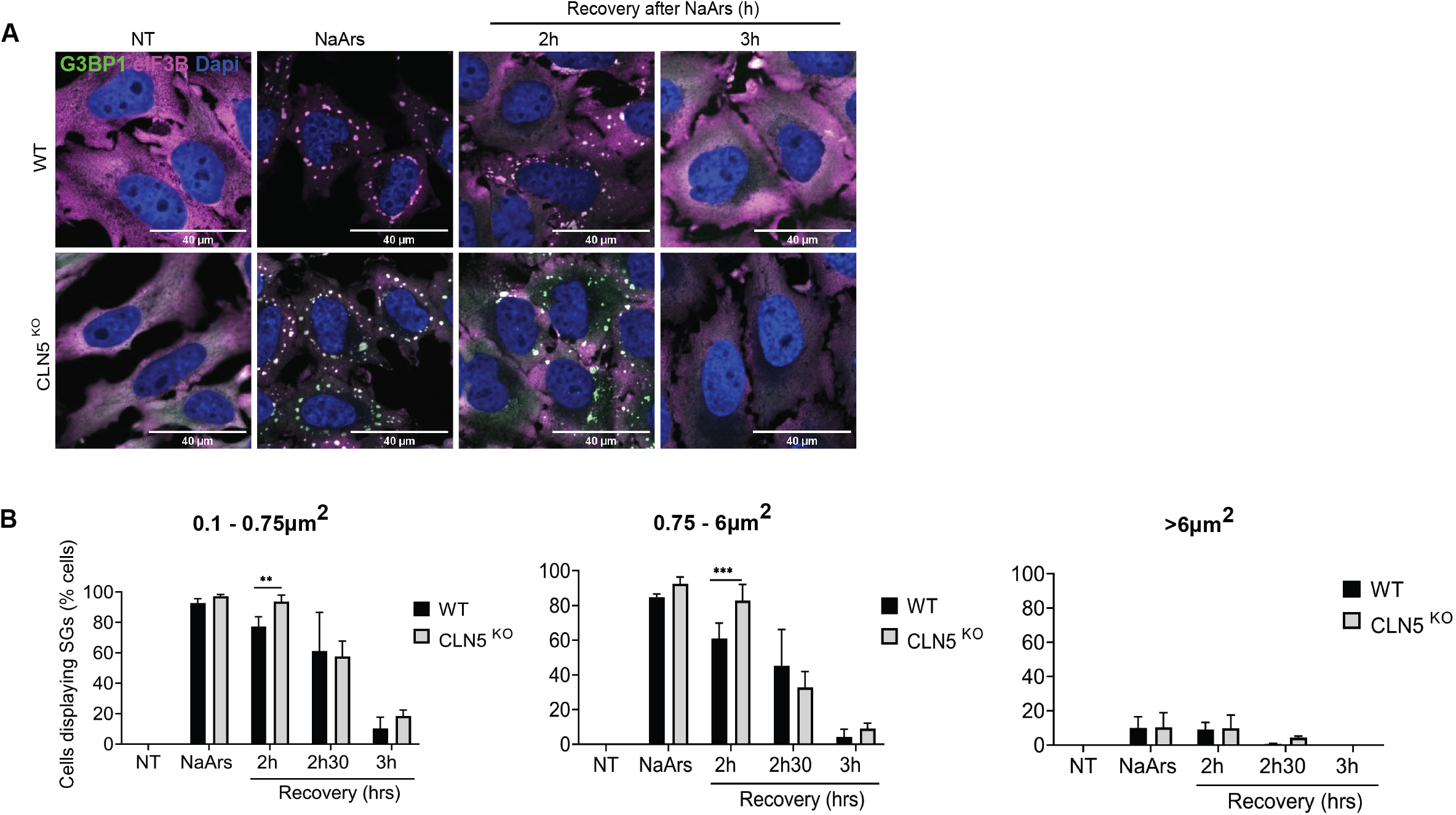
Impaired SG clearance is a CLN3 specific phenotype. (A) Rate of SG recovery was assessed in alternative BD model, CLN5^KO^ HeLa cells. Representative immunofluorescent images show time course of SG clearance after NaArs challenge. Scale bar = 40μm^2^. Cells stained for G3BP1 (Green) and eIF3B (Magenta). (B) Quantification of small (0.1-0.75 μm^2^), medium (0.75-6 μm^2^) and large (>6 μm^2^) shown as %Cells displaying SGs. Results shown mean ± SD, n=3. CLN5^KO^ HeLa cells display higher % of cells with small (P=0.03) and medium sized (P= 0.005) SGs at 2h post stress. Quantification performed using Image J plugin Aggrecount Statistical analysis was carried out using multiple t-tests.

### Delayed recovery of SGs in CLN3^KO^ cells is independent from eIF2α signalling or chaperones associated with SG disassembly

Following transient stress, the dissolution of SGs and translation re-entry follows the dephosphorylation of the α sub-unit of eIF2α (40). Chronic activation of the integrated stress response (ISR) via phosphorylated-eIF2α (p-eIF2α) is associated with SG persistence, toxicity and neuronal death (41). To assess whether the delayed SG clearance in CLN3^KO^ cells is linked to a defect in eIF2α signalling, p-eIF2α (S51) levels were measured in response to NaArs and at 3h following stress removal by western blot analysis. As expected, p-eIF2α (S51) levels increase significantly in WT HeLa cells upon NaArs induction and returned to baseline levels at 3h following NaArs removal (Fig 5A). NaArs stimulation in CLN3^KO^ HeLa cells resulted in similar level p-eIF2α activation which also returned to baseline levels by the end of the recovery period. These results suggest eIF2α is appropriately dephosphorylated at 3h post stress in CLN3^KO^ HeLa cells. Therefore, it is unlikely that the increased persistence of SGs observed in CLN3 deleted cells is linked to a defect in eIF2α signalling.

**Figure 5:**
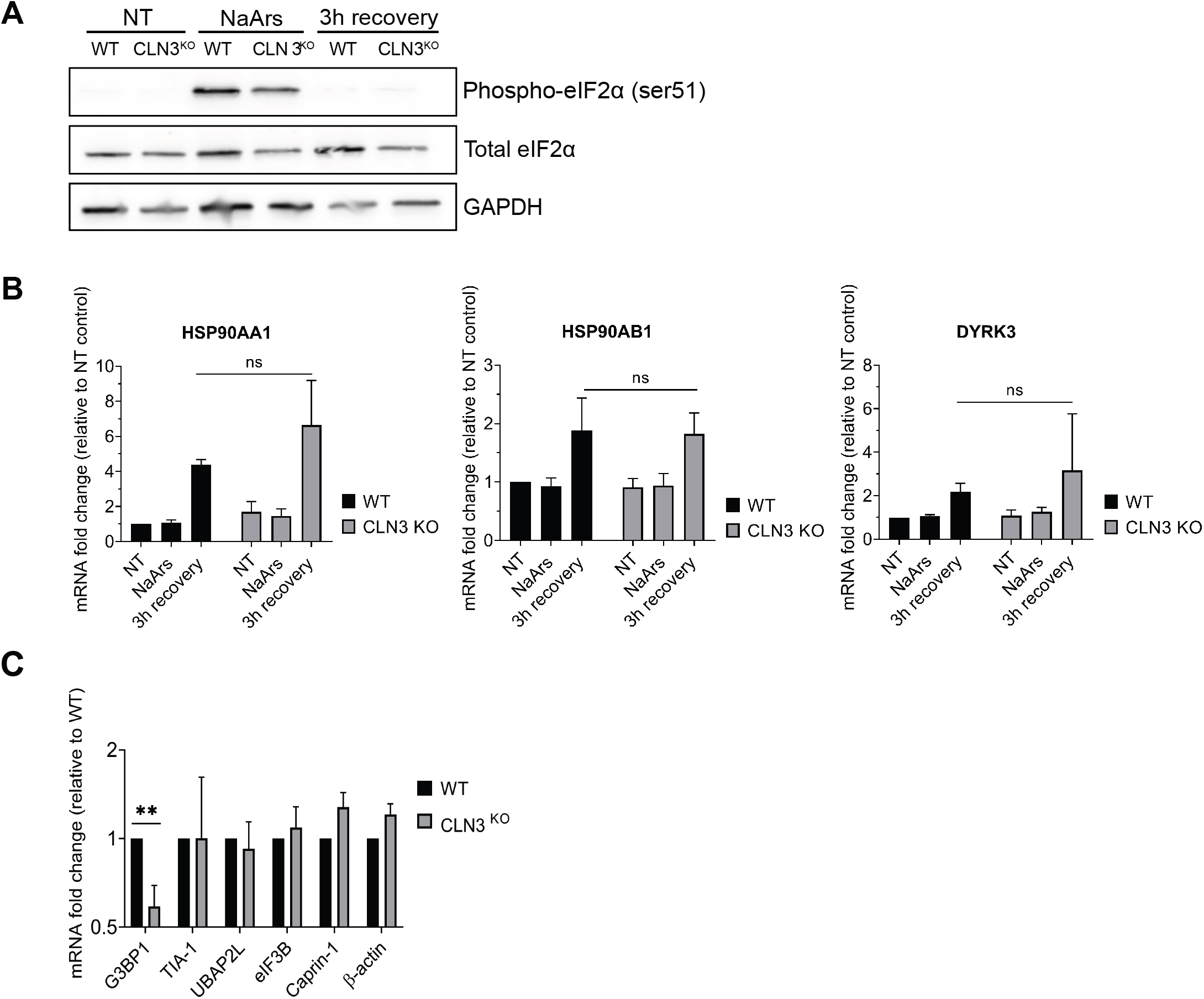
(A) Immunoblotting analysis of eIF2α phosphorylation (ser51) shows no difference in CLN3^KO^ HeLa cells compared to WT in NT, NaArs treated and 3h post NaArs conditions. (B) RT-qPCR analysis of SG chaperones HSP90AA1, HSP90AB1 and DYRK3 mRNA expression in response to NaArs challenge and 3h post stress. Results show mean ± SD, n=3, normalised to β-tubulin mRNA and shown relative to WT NT expression level. Statistical analysis performed using multiple t-test.

Upon stress, SGs sequester mTORC1 to modulate cell metabolism and growth to subsequently promote stress adaptation. Upon stress relief, HSP90, a molecular chaperone implicated in human disease, stabilizes DYRK3 which drives SG dissolution and the release of mTORC1, enabling restoration of translation and other mTORC1 dependent pathways (42-44). Aberrant signalling through this axis is associated with persistent SGs and ALS. To probe the impact of CLN3^KO^ on this pathway, we measured the mRNA expression of the stress inducible and constitutively expressed HSP90 subunits HSP90AA1 and HSP90AB, respectively, as well as the HSP90 substrate DYRK3 in non-treated, stressed and 3h post stress conditions (Fig5B). As expected, we observed that in WT cells the expression of HSP90AA1 and HSP90B, and DYRK3 were increased at the end of the recovery period from NaArs stimulation, corresponding to the kinetic of SG disassembly (Fig5B). A similar pattern was observed in CLN3^KO^ cells, with no significant difference in the chaperone induction between WT and CLN3^KO^ cells. Overall, this suggests that impaired recovery from stress and SG disassembly is not driven by a defect in chaperone activity.

### Loss of CLN3 is associated with reduced expression of the SG-scaffolding protein G3BP1, but not its ability to undergo SG-like phase transition

Appropriate expression and availability of key SG resident RBPs is a central determinant of SG size and function, with SG size governing functional interactions with other membrane-less organelles (45). Given that CLN3^KO^ cells display significantly less large SGs (>6μm2) after NaArs treatment, we asked if this was due to altered expression of key SG ‘node’ proteins. To answer this, we measured the expression of SG proteins G3BP1, TIA-1, UBAP2L, eIF3B and CAPRIN-1 in untreated conditions (Fig5C). While the mRNA expression level of most markers was unchanged between WT and CLN3^KO^ HeLa cells, we observed a two-fold reduction in the expression of G3BP1 in CLN3^KO^ when compared to WT HeLa cells.

It has been proposed that the assembly of SG is driven by the phase separation of G3BP1 once the percolation threshold has been reached (46). Furthermore, the ability of G3BP1 to undergo LLPS, and drive SG assembly in cells under stress is dependent upon its dynamic post-translational modifications, such as the removal of methyl groups on arginine residues and the phosphorylation status of serine 99 (47,48). These modifications prevent unnecessary and untimely formation of G3BP1 aggregates by neutralising the physical ability of G3BP1 to homo-and hetero-oligomerize with other SG-associated proteins and non-translating RNAs. Considering LLPS occurs in a concentration dependent manner, we asked how the reduced G3BP1 expression observed in CLN3^KO^ cells would affects its propensity for phase separation *in vitro* using biotin isoxazole (b-isox), a surrogate for SG formation (49). B-isox addition, in conjunction with an EDTA-EGTA lysis buffer releasing transcripts from polysomes, results *in vitro* in selective condensation of the low-complexity aggregation-prone proteins via phase separation (50). The b-isox-induced aggregates can then be fractionated from the soluble fraction by centrifugation, respectively defined as pellet and supernatant, and their protein contents analysed by immunoblotting (Fig6A). Quantification of immunoblotting analysis showed no difference in the levels of G3BP1 recovered in the b-isox precipitated fractions (pellets) compared to the inputs in the extracts from WT or CLN3^KO^ cells (Fig 6B). Moreover, canonical SG markers Caprin1 and UBAP2L were also precipitated with G3BP1, suggesting both the precipitation of low-complexity proteins and the interaction of G3BP1 with non-translating mRNPs. Overall, these data suggest the reduced level of G3BP1 associated with loss of CLN3 do not impact the ability of other SG proteins to phase separate and condense into SGs.

**Figure 6:**
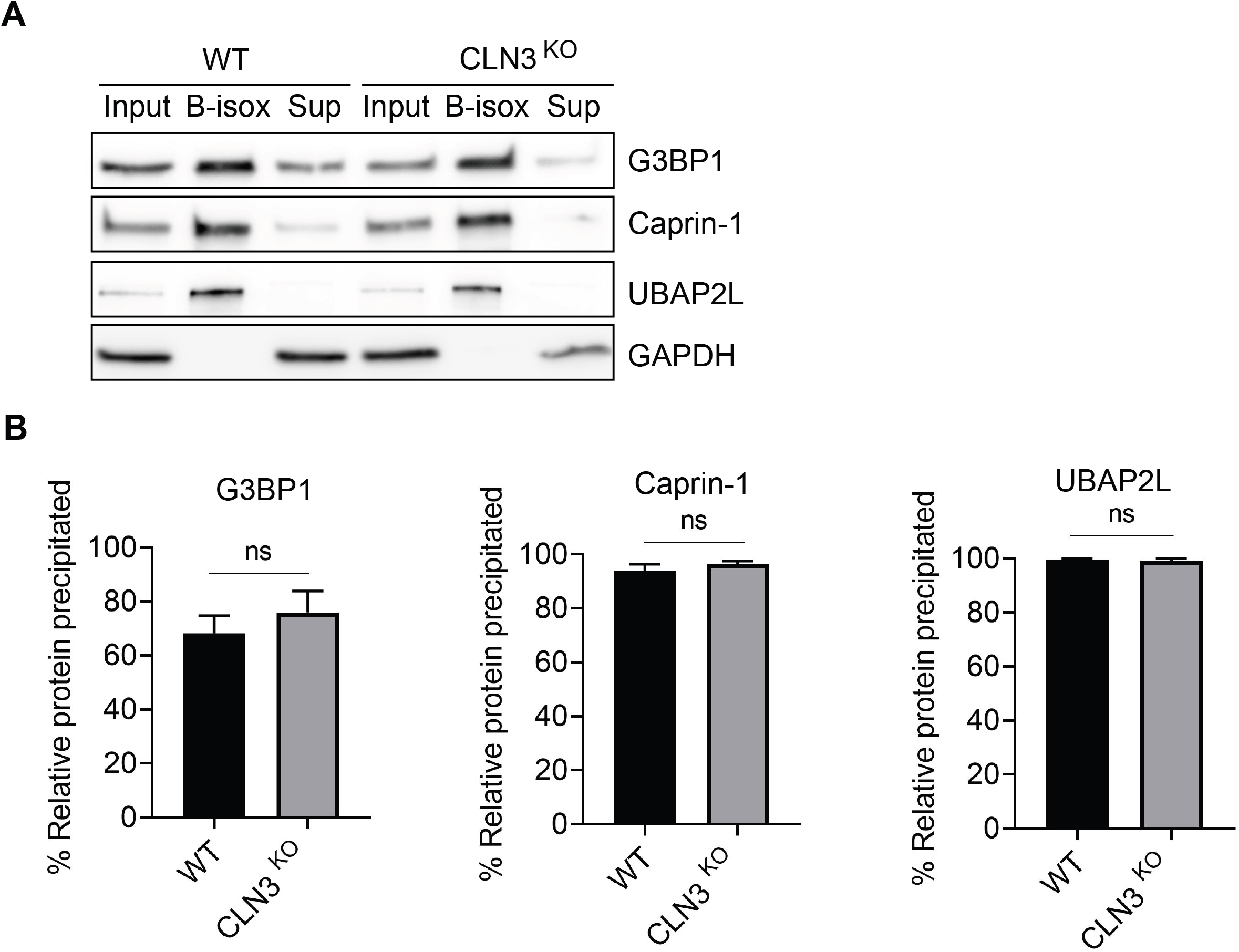
*In vitro* induced LLPS leads to SG-like aggregate formation in HeLa cells. (A) Analysis of key SG nodes mRNA expression by RT-qPCR shows significantly reduced G3BP1 expression in CLN3^KO^ HeLa cells. RT-qPCR results show mean ± SD, n=3, normalised to β-tubulin mRNA and shown relative to WT expression level. Statistical analysis was performed using unpaired t-test (** p=0.0025) (B) Western blot analysis of distribution of key SG nodes proteins in WT and CLN3^KO^ HeLa cells. For each genotype samples were loaded as input, b-isox pellet and b-isox supernatant from left to right. (C) Graphs show relative precipitation of SG nodes in b-isox pellet compared to input. Results shown as mean ± SD, n=3. All results non-significant upon analysis by unpaired t-test.

## Discussion

CLN3 batten disease displays multifactorial pathologies similar to those seen in age-associated neurodegenerative conditions, including metabolic and calcium signaling abnormalities (51), elevated reactive oxygen species (17), defective intracellular trafficking (13,39,52) and autophagy (14). We sought to investigate how loss of CLN3 modulated the SG pathway. We found that CLN3 KO cells have 1) altered cell metabolism and impaired protein translation 2) demonstrate delayed SG formation and 3) display severe delays in SG resolution.

We first confirmed our CLN3^KO^ HeLa cell model displayed features of homeostatic imbalance, by assessing rates of glycolysis and oxidative phosphorylation. These data support that CLN3^KO^ lowers the basal rates of both glycolysis and oxidative phosphorylation, and impairs the ability of cells to handle additional metabolic stress. Thus, our model recapitulates some of the pathological features associated with CLN3 batten disease. Furthermore, we detected perturbations in stress signaling pathways which are critical for adaptation to adverse cellular environments. We observed reduced phosphorylation and thus activation of the heat shock protein 27 (HSP27), JNK1/2/3 and p38α. HSP27 has numerous protective capacities and is implicated in many disease contexts including cancer, cardiovascular and neurodegenerative diseases where it plays a role in reducing ROS damage, inhibiting various cell death pathways and promoting protein folding (53). Similarly, mitogen-activated protein kinase (MAPK) signaling pathways, specifically JNK and p38α pathways are activated by environmental and genotoxic stressors and orchestrate multiple cellular defense mechanisms (54). JNK signaling is implicated in neuronal differentiation, motility, metabolism and apoptosis and abnormal signaling is linked to neuronal pathology including ALS and Alzheimer’s disease (55). In addition, previous studies using drosophila models have demonstrated a genetic interaction between CLN3 and the JNK signaling pathway (18). Together, our CLN3^KO^ HeLa cells display altered homeostasis at basal levels, sensitivity to additional metabolic stress and altered stress signaling which may suggest a role for CLN3 in the cellular stress response.

In response to stressful stimuli, SGs sequester non-translating mRNA, RBPs and signaling molecules to regulate a network of cellular pathways to regain homeostasis. Given we have observed metabolic, signaling and proteostasis defects and an increased sensitivity to additional stressors, we probed the ability of CLN3^KO^ cells to mount an SG response to exogenous stress. Cells were stimulated with sodium arsenite over a time-course and the frequency/number of SGs was quantified throughout the duration of stress, capturing SG heterogeneity by defining foci as either small (0.1-0.75μm^2^), medium (0.75-6 μm^2^) or large (>6 μm^2^). Surprisingly these set of experiments revealed that significantly fewer large SGs are present at the later stages of stress induction in CLN3^KO^ compared to WT HeLa cells. SG assembly occurs in a step-wise manner when translation initiation is stalled. Firstly, non-translating mRNP complexes oligomerise into stable core structures. Secondly, cores act as scaffolds for the recruitment of further mRNP complexes that undergo many fleeting and promiscuous interactions with one another resulting in the formation of the outer dynamic shell and phase separation from surrounding cytoplasm. Thirdly, during the latter stages of assembly, small SGs undergo multiple fusion events to form larger, mature SGs that contain multiple cores (56). The reduction in large SGs observed during the later stages of assembly may be due to impairments in the fusion of small SGs that form early in the stress response. Small assemblies can be directly or indirectly transported along microtubules during cell transport. Studies have shown that stress induced RNA granules can be tethered to motile lysosomes for long distance travel (57). One possibility is that defective lysosome function and localisation (8) in CLN3 batten disease impairs lysosome motility which directly effects the ability of RNA granule to ‘hitchhike’ during cell transport, in turn preventing fusion events that lead to granule maturation. Future experiments should aim to further investigate this interaction between SG dynamics and lysosomes.

To further probe the molecular mechanism that could explain the defect in SG assembly, we assessed the levels of several SG markers in CLN3^KO^ cells. This revealed that the key SG nucleating factor G3BP1 was downregulated by 2-fold at the mRNA level in CLN3^KO^ cells, while other transcripts encoding proteins important within the SG core were consistent between genotypes. G3BP1 encodes a multifunctional RBP that interacts with multiple cell pathways including stress response and RNA metabolism, suggesting it’s importance for maintaining homeostasis (58). In SG formation, G3BP1 is thought to be a critical regulator of LLPS due to its high valency for RNA binding (23). Reduced G3BP1 mRNA is a feature of ALS with TDP-43 pathology (59). Importantly, despite the reduction in transcript level, G3BP1 protein level and its ability to undergo phase separation was unimpaired in CLN3^KO^ cells. This may suggest a complex pattern of translational control for G3BP1 mRNA associated with CLN3. Interestingly previous work identified SG-dependent and independent roles for G3BP1 in promotion transcript-selective translation and guiding transcript partitioning to reprogram mRNA translation and support stress adaptation (60).

SG dynamics are influenced by a tightly regulated network of protein quality control (PQC) pathways that maintain protein homeostasis by promoting the timely disassembly of SGs after stress. Recent work on SG disassembly has shown it is a highly context specific process-with stressor type and duration influencing the mechanism of clearance (61,62). Failure of PQC pathways is associated with aging and disease and leads to persistent SGs, a putative cause of neurodegeneration (63). Given the observed impairment in SG assembly, we sought to investigate how CLN3^KO^ impacted the SG disassembly process after arsenite challenge. Our data showed an increased persistence of small and medium SGs after stress release associated with CLN3^KO^ cells, which could not simply be explained by prolonged eIF2α activation.

SG integrity is also maintained by a network of molecular chaperones that prevent protein misfolding within the granule and aid the recruitment of other essential SG factors. Heat shock protein 90 (HSP90) is a major chaperone implicated in SG dissolution and stress adaptation. By modulating the activity of dual-specificity tyrosine-phosphorylation-regulated kinase 3 (DYRK3), HSP90 couples SG dissolution with the release of MTORC1 and re-initiation of metabolism. Reduced activity and expression of DYRK3 is seen in ALS and is causative of SG clearance defects and decreased viability (44). Although we observed an increase in expression of HSP90 subunits during the 3 h recovery phase, there was no difference in the change of expression between WT and CLN3^KO^ cells, suggesting that in our model HSP90 modulation of DYRK3 is intact, and any impairment in SG clearance is not due to inactivation of DYRK3. Recent work has shed light on the role of post translational modifications (PTMs) in condensate assembly (23,64). In particular, for the key SG node G3BP1, phosphorylation at residues S149 and S232 flanking its acidic IDR have been proposed to increase the threshold for phase separation likely keeping G3BP1 in an SG-incompetent conformation (22,23). Additionally, the G3BP1 RGG domain is the target of differential arginine methylation by the protein arginine methyltransferases PRMT1 and PRMT5, with methylation at residues R435, R447 and R460 strongly inhibiting SG formation, while loss of PRMT1 and PRMT5 activity is associated with a small SG phenotype (47). Together, PTMs of G3BP1 represent a way of fine-tuning SG assembly and future studies should aim at their investigating in the context of CLN3 absence or presence.

The equilibrium of cellular protein levels (termed proteostasis) is maintained by a network of interactions between numerous biochemical pathways. Protein synthesis, folding, post-translational modification and degradation are highly regulated to ensure the functional requirement of cells are met while ensuring PQC and degradation pathways are not overwhelmed (65,66). Exposure to stress can perturb the delicate balance of interactions that maintain proteostasis, which may subsequently lead to pathological compensatory changes. Equally, reduced efficiency of degradation pathways by disease associated mutations or aging can result in the accumulation of toxic protein aggregates (67). Using the ribopuromycylation assay to measure translation at single cell level, we observed a significantly reduced rate of translation under basal conditions in CLN3^KO^ cells. These results were further confirmed by polysome profiling fractionation experiments demonstrating a reduced association of mRNAs with highly translating ribosomes, and a reduced polysome/monosome ratio in CLN3^KO^ cells, overall suggesting there are fewer mRNA-bound ribosomes in steady state conditions and impaired translational activity associated with loss of CLN3. Recent work in a yeast model has shown a genetic interaction between the CLN3 homologue, btn1, and genes encoding ribosomal proteins suggesting a link between CLN3 function and protein synthesis (68). Work in the same HeLa KO system used in this study has also shown CLN3^KO^ impairs the late stages of autophagy (13,14). Autophagy is one of the major catabolic routes of protein degradation and essential for cell viability. Impairment in the clearance of proteins disturbs the balance of protein homeostasis and cells may establish compensatory mechanisms to restore equilibrium. It is plausible that in CLN3^KO^ cells, global translation is reduced to lessen the burden on deficient the autophagy pathway. Data from the literature further support this, as previous work has shown a direct and functional interaction between CLN3 and Rab7a, a small GTP-ase implicated in the function of late endosomes (13). Late endosomal Rab7a has been shown to bear mRNA that is translated by associated ribosomes and translational machinery. Further these complexes of endosomes and RNP granules dock onto mitochondria and translate mRNA essential for mitochondrial vitality (69). Mutations causative of CLN3 batten disease disturb the distribution of late endosomal and lysosomal compartments resulting in perinuclear clustering and impaired motility (8). We suggest that the physical interaction between late endosome/ lysosome and translating RNP granules may be impaired in CLN3^KO^ cells. Furthermore, decreased site-specific translation of mitochondrial proteins which is dependent on this interaction would likely contribute to impaired mitochondrial activity. Together, these data suggest that the requirement for CLN3 in translation is conserved between yeast and mammalian cells. It remains unanswered what mRNAs are being translationally repressed, whether it is a global phenomenon or specific to certain populations of transcripts. Future work expanding on the polysome fractionation performed in this study may shed light on this question and unravel the deregulated pathways that contribute towards CLN3 disease pathogenesis.

This study has shown for the first time a potential interaction between CLN3 pathology and SG assembly, disassembly, and the regulation of steady-state translation. Given the role of SGs in the reorganisation of cellular contents to maintain homeostasis in steady-state conditions and in response to stressful stimuli, and recent work highlighting the importance of lysosomes in SG shuttling in neurons to avoid neurodegeneration, our study paves the way for further investigation in patient-derived cells to explore the link between lysosomes, SG dynamics and modulation in CLN3 disease.

## Experimental Procedures

### Cell culture

HeLa WT, CLN3 and CLN5 cell lines were maintained in Dulbecco’s modified Eagle’s medium (DMEM) supplemented with 100 U/ml penicillin, 100 g/ml streptomycin and 10% FBS. at 37°C in a humidified chamber at 95% air and 5% CO2. CLN3^KO^, CLN3^KO^ and Rab7A^KO^ HeLa cells have been described previously (13,39,70). Cells were seeded at a density of 5×10^6^/ well in 24 well plates, 2×10^6^/ well in 6 well plates and 10×10^6^/well in 15cm dishes.

### Stress induction and recovery assays

For SG induction assay, stress was induced with 0.5mM sodium arsenite (NaArs, Sigma) for 5, 10, 20 or 45 min at 37°C. For the SG recovery assay, granules were induced with the addition of 0.5mM NaArs for 45 min. NaArs was then washed out with fresh DMEM for 5 min at 37°C and this process repeated 3 times. Fresh media was added for the final time and cells allowed to recover for a maximum of 3 hours. Immunofluorescence staining and image acquisition was carried out as indicated below. All image were processed and analysed using Image J software package Fiji (http://fiji.sc/wiki/index.php/Fiji) and plugin Aggrecount (https://aggrecount.github.io) was used to quantify SGs in an automated and non-biased manner (37). Graphpad prism was used for data presentation and analysis was performed using multiple t-tests.

### Ribopuromycyclation (RPM) assay

Quantification of *de novo* protein synthesis was performed as described in (53). To capture translation efficacy in non-treated (baseline) and stressed condition, cells were treated with 0.5mM NaArs for 45 mins. Then, prior to fixation, cells were treated with 10μg/ml of puromycin (Sigma) for 5 min at 37°C to label the nascent polypeptide chains before addition of 180μM of emetine (Sigma) to block the translation elongation with a further incubation of 2 min at RT. Cells were rinsed with warm DMEM then fixed with 200μl 4% PFA for 20 min at RT and washed 3 times in PBS. Puromycin incorporation was quantified using Image J software package Fiji. The raw integrated density values were recorded for ∼100 cells per condition. Graphpad prism was used for data presentation and analysis was performed using Mann Whitney U test.

### Polysomes fractionation

Cells were seeded to 10cm^2^ plates. Upon reaching ∼80% confluency, cells were treated with 100μg/mL of cycloheximide (CHX-Sigma) for 3min at 37°C to stall translating ribosomes onto the transcripts before being placed on ice, washed twice with cold PBS(-) (Invitrogen) containing 100μg/ml CHX, then scraped into 15ml falcon tubes, pelleted by centrifugation snap frozen in liquid nitrogen and stored at −80°C. To separate polysomes, samples were layered onto a 10–50% sucrose gradient in lysis buffer and centrifuged in an SW40Ti rotor (Beckman Coulter, High Wycombe, UK) at 38,000 rpm for 2 h. Gradients were fractionated into 1-ml fractions using a FoxyR1 collection system (Teledyne ISCO, Lincoln NE), and UV absorbance was monitored at 254 nm. To induce run-off of polysomes, cycloheximide was omitted from the lysis and gradient buffers and replaced with 10 mM EDTA.

### Biotin isoxazole (B-isox) precipitation

Protocol was performed as described in (52). Briefly, cells were seeded to 10cm^2^ plates at 2×10^6 cells per plate 24 hours prior to experiment. Cells were placed on ice and washed twice with cold PBS (from 10x solution–Lonza). Whilst in PBS, cells were gently disassociated using a cell scraper and transferred to15ml falcon tube and centrifuged at 2000 RPM RT for 3 min. The supernatant was removed, and pellet was snap frozen in liquid nitrogen. Cells were lysed on ice in cold EE buffer (Hepes pH 7.4 50mM, NaCl 200mM, Igepal 0.1%, EDTA pH8 1mM, EGTA pH8 2.5mM, Glycerol 10%, DTT 1μM supplemented with RNAsin (Promega). Lysates were transferred to 1.5ml tubes and incubated with agitation for 20mins at 4°C then centrifuged at 13,000 RPM 15 min at 4°C. 50μl of the supernatant was kept as “input”, mixed with an equal volume of 2x Loading buffer (Cell Signalling) and boiled at 95°C for 5min. The remaining supernatant was supplemented with 100μM of b-isox (Sigma) and the precipitation reactions were carried out for 90min at 4°C with agitation. 50μl supernatant was kept as “soluble fraction” and mixed in equal volume with 2x loading buffer and boiled 95°C for 5min. The pellet containing precipitated aggregates was rinsed twice in cold EE buffer and spun at 10,000RPM 10 min at 4°C then resuspended in 200μl x1 loading buffer as “insoluble fraction” and boiled at 95°C for 5min. Fractions were further analysed by western blot performed as indicated. Quantification was performed by calculating relative precipitation of protein in b-isox pellet compared to input. Data was analysed in Graphpad prism and statistics performed using unpaired t-test.

### Phospho-kinase Arrays

Cells were seeded to 15cm^2^ plates at 5×10^6 cells per plate 48 hours prior to experiment. Prior to harvesting, cells were treated with 0.5mM NaArs for 45 min and harvested straight away by scraping into 15ml tube, followed by centrifugation and snap freezing the pellet or washed 3 times in DMEM to remove NaArs and incubated for further 3 hours before harvesting. Cell pellets were lysed in 1ml lysis buffer from Human phospho-kinase array kit (R&D systems, ARY003C) supplemented with protease inhibitor cocktail (Roche). Protein concentration was measured with BCA pierce kit (Fisher) and 400μg protein used for each analysis. Arrays were performed as per manufacturers guidelines. Images were acquired using the VILBER imaging system and quantified in FIJI. Data presentation and analysis was performed in Graphpad prism using multiple t-tests.

### Immunoblotting

Cells were plated at 2×10^5^ in 6 well plates. At the indicated times, cells were lysed in 150μL of 1x Gel Loading Buffer (New England Biolabs), sonicated and boiled 5min at 95°C. Cell lysates were separated by SDS-PAGE and protein was transferred to nitrocellulose or polyvinylidene difluoride membranes. These were then probed with the following primary antibodies: rabbit anti-eIF2a (1:1,000, Cell Signalling), mouse anti-P-eIF2α (1:1,000, Cell signalling), rabbit anti-G3BP1 (1:2,000, Sigma), rabbit anti-Caprin1 (1:1,000, Bethyl Laboratories), goat anti-eIF3B (1:2,000, Santa Cruz). mouse anti-GAPDH (1:20,000, Invitrogen); followed by incubation with the appropriate peroxidase labelled secondary antibodies (Dako) and chemiluminescence development using the Clarity Western ECL Substrate (Bio-Rad). The results were acquired using the VILBER imaging system.

### Immunofluorescence

5×10^5^ Hela cells were plated on glass coverslips in 24 well plate and fixed with 4% PFA (sigma) in PBS for 20 min at RT, washed in PBS and stored at 4°C. Cells were permeabilised with 0.1% Triton-×100 (Sigma) in PBS for 7 mins then blocked with 0.5% BSA (Fisher) in PBS for 1h. Coverslips were then incubated with 200μl of primary antibody solution for 1 h at RT and washed 3 times with PBS before incubation with secondary antibody solution containing 0,1μg/mL DAPI solution (Sigma) for 1 h at RT. After 3 washes with PBS, coverslips were mounted to microscope slides with 7μl Mowiol 488 (Sigma). Confocal microscopy was performed on a Ti-Eclipse -A1MP Multiphoton Confocal Microscope (Nikon) using the Nikon acquisition software NIS-Elements AR. Primary antibody dilution: Rabbit anti-G3BP1 (1:600, Sigma), Goat anti-eIF3B (1:600, Santa Cruz), mouse anti puromycin (1:5, http://dshb.biology.uiowa.edu/PMY-2A4), Rabbit anti-UBAP2L (1:600, manufacturer). Secondary antibodies were all purchased from Invitrogen: Chicken anti-mouse Alexa 488, donkey anti-goat Alexa 555 and goat anti-rabbit 488.

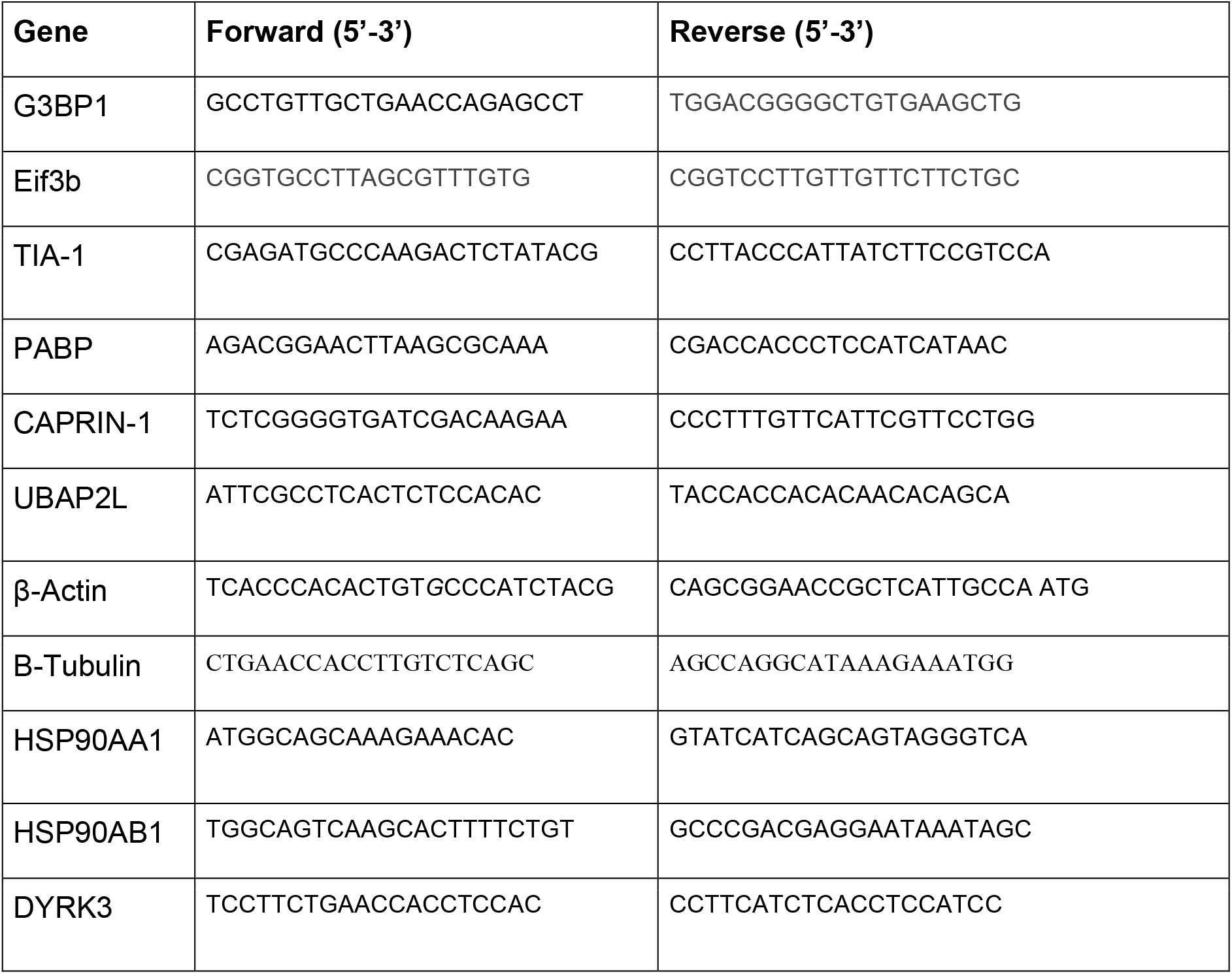

### RT-qPCR

Prior to experiment cells were seeded at a density of 2×10^5^ cells/ well 6 well plates. Cells were washed in cold PBS then total RNA isolated using quick-RNA extraction mini prep kit (Zymo research) according to the manufacturer’s instructions before quantification on Nanodrop. Reverse transcriptions were carried out on 0.5 μg of purified RNA using the Precision nanoScript2 Reverse Transcription kit (Primer Design) and qPCR was performed on 5μL of the cDNA libraries diluted at 1:5 using the Precision Plus 2x 598 qPCR Mastermix (Primer Design) according to the manufacturer’s instructions and the Quant Studio 7 Flex (Applied Biosystems).

### Primer list

#### Seahorse Methods

The measurement of rates of oxidative phosphorylation and glycolysis in wild type HeLa cells and CLN3 KO HeLa cells was carried out using the Agilent Seahorse XF Cell Energy Phenotype Kit (Agilent #103325-100) using the Agilent Seahorse XFe 96 Analyzer, according to the manufactures instructions. WT or CLN3 KO HeLa cells were plated into the Seahorse XF Cell Culture Microplate at a concentration of 20,000 cells/well in DMEM (Gibco #41966-029) + 10% FBS (Sigma #F9665). Seahorse XF Assay media was supplemented with 1 mM pyruvate, 2 mM glutamine, and 10 mM glucose (Agilent # 103681-100). Oligomycin and FCCP were reconstituted to concentrations of 10 μM and 5 μM, respectively. The metabolic phenotype was measured using the Cell Energy Phenotype assay template. Data was normalised by protein concentration using the Protein Assay kit (Biorad #5000116). GraphPad Prism was used for data presentation and statistical analysis using the unpaired t-test.

## Data availability

All the data are included in the article. All unique reagents generated in this study are available from the corresponding authors with a completed Materials Transfer Agreement.

## Supporting information

This article contains supporting information

## Acknowledgements

We thank M. Brocard for helpful discussions and technical assistance during the early stages of this study.

## Author contributions

N. L., P. J. M. and S. L. conceptualization; N. L., E. L. R. and P. J. M. methodology; N. L., E. L. R., D. N., and P. J. M. formal analysis; N. L., E. L. R., S. Y., A. K., and N. J. R. investigation; N. L. and E. L. R. writing–original draft; E. L. R., N. L., G. H., S. L. and P. J. M. writing–review & editing; N.L. supervision.

## Funding and additional information

Work in N.L.’s laboratory is supported by Medical Research Council and the European Union Joint Programme in Neurodegenerative Diseases grant [grant number MR/R02426X/1]. P.J.M’s laboratory is supported by the Medical Research Council and the European Union Joint Programme in Neurodegenerative Diseases grant MR/R02426X/1] and the NIH-JPND grant MR/S022465/1] and N.J.R is part of an MRC-DTP. Work in S.L.’s laboratory is supported by grants from the Canadian Institutes of Health Research (Joint Programme in Neurodegenerative Diseases (Neuronode) [grant number ENG-155186] and Project grant [grant number PJT-173419], the Canadian Foundation for Innovation [grant number 35258], and ForeBatten Foundation. Work in G.H.’s laboratory is supported by grants from the Deutsche Forschungsgemeinschaft [grant number HE 3220/4-1] and the NCL Foundation.

## Figures Legends

**Supporting figure S1:**
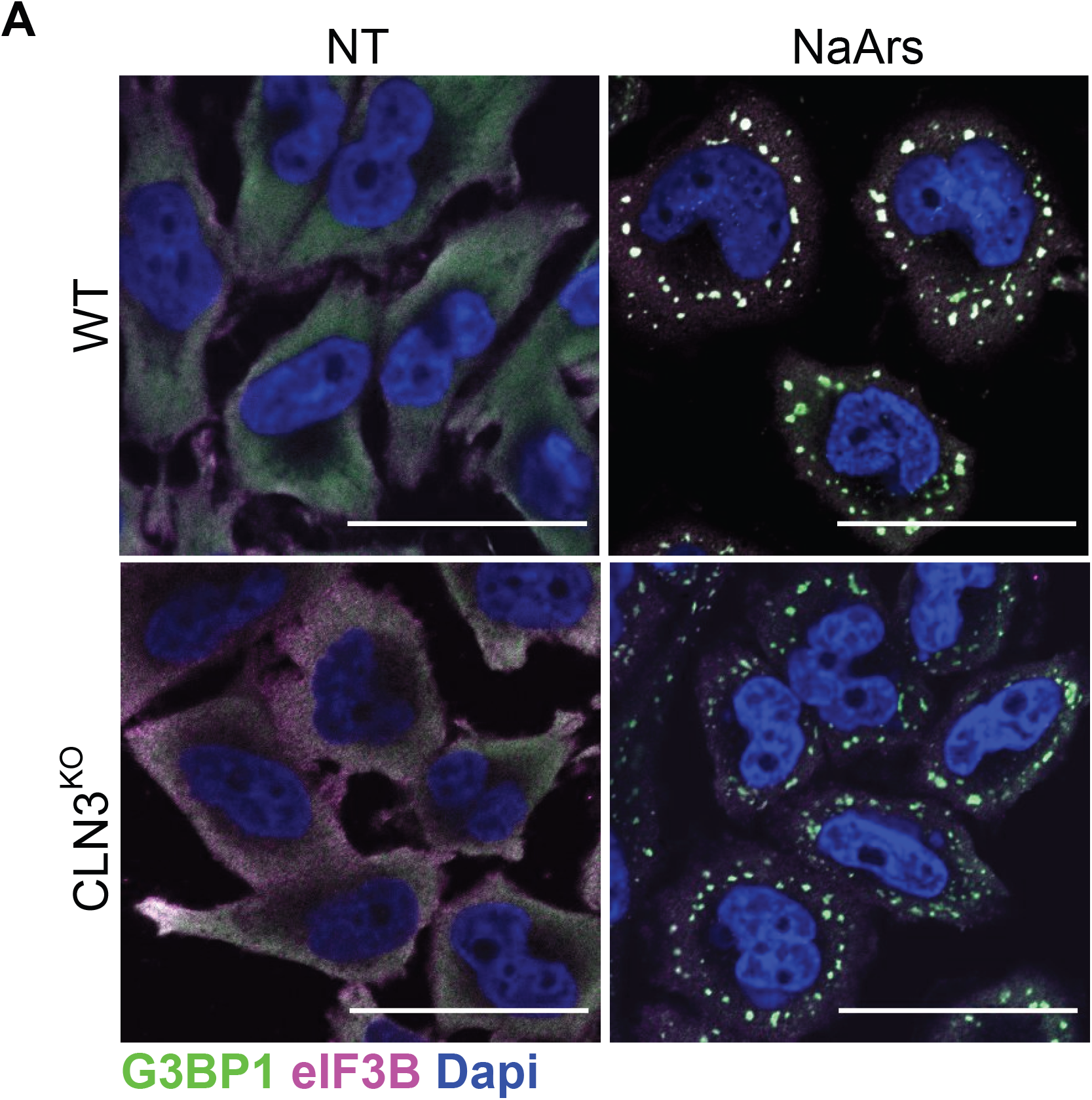
(A) Representative immunofluorescent images show colocalisation of SG markers G3BP1 (green) and eIF3B (magenta) at 20mins into NaArs induction time course. Scale bar = 40 μm^2.^

**Supporting figure S2:**
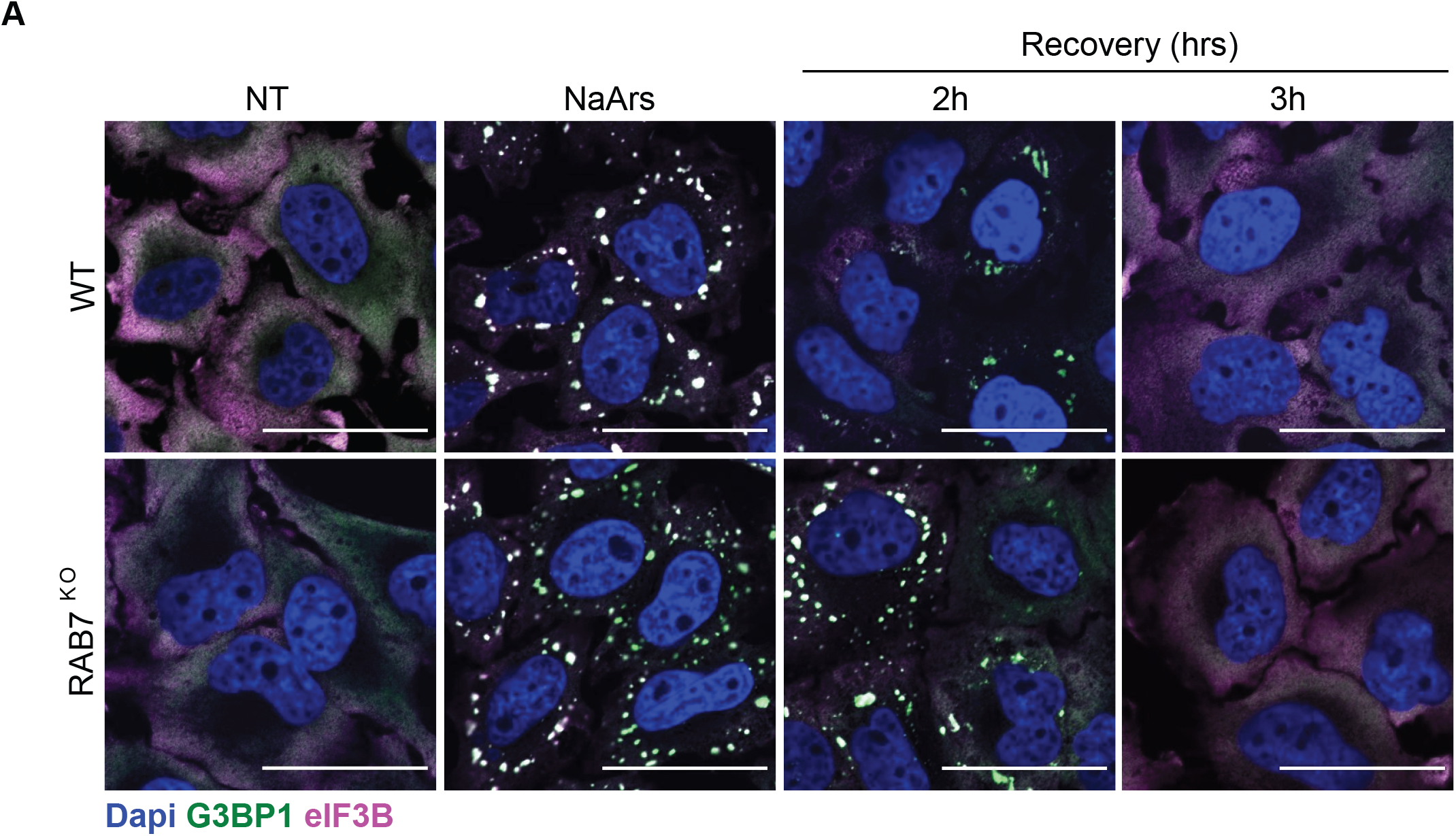
SG clearance is unaffected by loss of Rab7A. (A) Representative immunofluorescent images stained for G3BP1 (green), eIF3B (magenta) and nuclear stain, dapi (blue). Scale bar = 40 μm^2^.

